# New resistance to bacterial wilt in heat-stressed tomato is revealed by two-reference Genome Wide Association

**DOI:** 10.1101/2025.07.28.667131

**Authors:** Adrien Belny, Henri Desaint, Mylène Rigal, Sébastien Carrère, Jérôme Gouzy, Gregori Bonnet, Céline Labourey, Elise Albert, Laurent Grivet, Fabrice Roux, Fabienne Vailleau, Tristan Mary-Huard, Richard Berthomé

## Abstract

Bacterial wilt, caused by bacterial strains of the *Ralstonia solanacearum* species complex, is one of the most harmful diseases striking many crops including tomato. Its spread is dependent upon temperature and humidity, which are expected to fluctuate strongly due to climate change. Previous results have highlighted that temperature elevation led to an increase in disease severity in commercial cultivars, whose resistance is quantitative and mostly relies on the Quantitative Trait Loci (*QTL*) *bwr-6* and *bwr-12*. In this study, we focused on temperature-dependent quantitative disease resistance (QDR) to bacterial wilt with the aim to unravel new resistance mechanisms that remain efficient at higher temperatures. For this purpose, a new panel of 189 accessions composed of tomato wild relatives, was assembled and sequenced thus creating a unique genomic resource. Its response to the *Ralstonia pseudosolanacearum* strain GMI1000 from three- to ten-days post-inoculation at 28°C and 32°C was explored. To discover the genetic basis underlying the responses of the panel, Genome-Wide Association (GWA) studies were conducted using the disease symptom scores recorded daily and monitored throughout the kinetics of the infection. To improve *QTL* detection, we have proposed a new approach using two reference genomes from within the panel. By correcting part of a single reference genome, especially when the only reference genome is a cultivar, this approach may be considered an alternative to pangenomic studies. As panel sequencing was highly resolutive, *QTL* positions allowed the identification of 44 candidate genes, which seemed to follow a temporal dynamic of activation after pathogen inoculation. Interestingly, no candidate genes were found to be common between the two phenotyping temperatures, highlighting the importance of the experimental design in addressing this type of question. Most of our quantitative disease resistance candidate genes belong to gene families described as being involved in immunity. Moreover, a significant proportion appears to be expressed in roots where bacterial infection occurs. Among them, two candidates are closely linked to the genomic positions of the *bwr-6* and *bwr-12 QTLs*, the main *QTLs* of bacterial wilt Quantitative Disease Resistance (QDR) studied whose mechanisms of action are still unknown.

**Author summary:** Bacterial wilt is a plant disease that affects more than 200 crop species (including tomato, potato and banana) leading to high yield losses. The most efficient way to deal with this disease is still the use of genetic resistance. However, previous studies have shown that numerous sources of resistance are negatively affected when plants face heat stress, which is alarming in a context of global warming. In this study, we have developed a new approach for discovering candidate genes in tomato capable of conferring thermostable resistance to bacterial wilt. Forty-four genes were sequentially detected over time, reflecting different temporal dynamics of induction after inoculation. Even if none were found to be common between the two temperatures, available information on their transcriptional regulation in roots and their involvement in immune processes confirms their relevance. Finally, we provide a short list of the candidate genes identified, some of which are currently undergoing functional validation and will be used in breeding programs to help overcome epidemics in future years.

## Introduction

Bacteria belonging to the *Ralstonia solanacearum* species complex (RSSC) are telluric phytopathogens that infect plant roots, colonizing and multiplying in the xylem, clogging vessels and leading to characteristic symptoms of plant wilting (1). The RSSC strains originate from tropical and sub-tropical regions and are organized into four major genetic clusters, called phylotypes. Interestingly, the geographical origins of the strains are correlated with their phylotype (2). Strains of the RSSC are able to infect a wide range of plant species from 53 botanical families and are responsible for various bacterial diseases that can lead to considerable economic losses in major crops (tomato, potato, eggplant and other Solanaceaous species), crops of local importance (banana, peanuts, etc), and ornamentals (pelargonium, rose) (3, 2).

To limit the occurrence of bacterial wilt and overcome disease outbreaks, integrated pest management combines multiple strategies (4). Promising developments in control methods are based on, or use, living organisms (5,6) and can complement traditional cultivation practices, such as crop rotation (7) and pesticide use (4). However, multiple studies have shown that genetic solutions remain the most efficient, cost-effective and eco-friendly means of tackling these diseases (8, 9).

The genetic determinism of plant resistance to pathogens has been extensively studied for many years, providing an interesting framework for improved understanding of plant- pathogen interactions and qualitative plant resistance, also known as gene-for-gene resistance. It has also resulted in a highly concise definition of plant immunity, reduced to the induction of a two-level response described in the well-known zig-zag model (10). Its mechanisms are based on the type of host receptors involved: PAMP-triggered immunity (PTI) or effector- triggered immunity (ETI). Results from the last two decades have brought to light the fact that mechanisms involved in plant immunity appear to share interconnected actors (11). Indeed, knowledge acquired from Quantitative Disease Resistance (QDR) studies (12, 13) revealed their polygenic nature. Moreover, QDR were sometimes found to confer resistance to multiple pathogens and to be modulated spatio-temporally. Thus, the simple dichotomic description of PTI versus ETI is no longer relevant. In contrast, knowledge of the genetic basis of plant defense responses to RSSC remain sparse. The best characterized resistance to *Ralstonia solacearum*, identified in the model plant *Arabidopsis thaliana,* is mediated by the nucleotide- binding, leucine-rich repeat (NLR) immune receptor pair RPS4 / RRS1 (14, 15, 16). Initially defined as a qualitative type of resistance, subsequent work demonstrated that this immunoreceptor pair conferred resistance to different strains of *R. solanacearum* (17, 14) as well as to other pathogenic species, such as *Colletotrichum higginsianum* or *Pseudomonas syringae* pv tomato DC3000 (18). Interestingly, two studies also demonstrated the role of RPS4 / RRS1 in QDR against *Xanthomonas campestris* pv*. campestris* strain CFBP6943 and *R. solanacearum* strain GMI1000 at 27°C (19, 20). Indeed, resistance to RSSC appears to be quantitative in most species studied. Evidence comes from studies of the repertoire of type III effectors, which are major determinants of the pathogenicity of RSSC strains, and genetic analyses of host resistance. These studies corroborate the fact that plant resistance to RSSC is strain-specific and mostly polygenic (8, 9). Overall, more than forty *QTLs* underlying plant defense responses to different strains of RSSC have been identified in *A. thaliana*, tobacco, potato, peanut, pepper, eggplant and tomato (21, 9), mainly by traditional *QTL* mapping approaches or, more recently, by exploring species genetic diversity and GWAS. However, the mechanisms involved remain unknown in most cases; except in few recent examples in *A. thaliana* in which candidate genes have been functionally validated. These candidates participate in resistance or susceptibility to the wild-type *R. solanacearum* strain GMI1000 or to four T3E mutants. (20, 22, 23).

Tomato (*Solanum lycopersicum*) constitutes one of the main Solanaceous species cultivated worldwide and serves as a model for fruit-bearing crops. As natural hosts of *R. solanacearum*, most tomato cultivars have been described to be susceptible to a wide range of strains belonging to the four phylotypes of the RSSC (24). Interestingly, *Solanum pimpinellifolium*, a wild relative of cultivated tomato, has contributed to the identification of resistance to over 45 different diseases (25, 26) including, for instance, resistance to *Phytophthora infestans* (27, 28), *Fusarium oxysporum* (I2 gene) (29), gray leaf spot disease (*Sm* gene) (30) and bacterial speck (*Pto* gene) (31). For bacterial wilt, the *S. lycopersicum* ‘Hawaii 7996’ cultivar is still the most tolerant to several strains of *R. solanacearum* (32) and is currently used in many breeding programs. Interestingly, the tolerance of this cultivar originates from *S. pimpinellifolium* accession PI127805A and is associated with at least seven *QTLs*, of which *bwr-6* and *bwr-12* are well studied (33). However, none of the genes underlying these *QTL* have been cloned and the genetic architecture of this QDR appears to be complex. Indeed, *bwr- 6* and *bwr-12* are made of several smaller *QTLs*: four for *bwr-6* and three for *bwr-12* (34).

A growing number of studies have assessed various combinations of plant-pathogen interactions under abiotic stress conditions (cold, high temperature, low nitrogen, drought) (35, 36), showing different plant defense responses to be strongly and negatively modulated by abiotic constraints. However the underlying mechanisms remain poorly understood, most studies being descriptive and supported by phenotypic data and transcriptomic data when analyzed (35, 37). For temperature increases, one of the climatic parameters predicted to fluctuate the most by the end of the century, a meta-analysis showed that 55% of a total of 145 genetic sources of resistance are negatively modulated by heat stress, which leads to increased disease severity (37). For RSSC, heat stress unveils the thermosensitivity of the *A. thaliana* RPS4/RRS1-R mediated resistance (20). This is also the case in Solanaceous species (38, 39). Indeed, increased soil temperatures can impair the bacterial wilt QDR for some tomato varieties (40). In Taiwan and other parts of South-East Asia, varieties with *bwr-12*-based QDR tend to be more susceptible to the bacteria at high temperatures. When phenotyped at elevated temperature, the ‘Hawaii 7996’ cultivar presented a wilting rate of between 28 % and 52 % depending on the trial (32). To date the Hawaii 7996 cultivar QDR remains the most reliable. However, new *QTL* conferring resilience to resistances to bacterial wilt in fluctuating environments have yet to be identified and characterized.

The natural diversity in the response of plant species to pathogens and genome-wide association studies (GWAS) has demonstrated their effectiveness in detecting *QTL* underlying plant defense responses (41, 42). However, the development of such strategies using accessions challenged by combined stresses is still scarce. Tomato wild-relatives originated from tropical and equatorial regions and evolved under more extreme climates, probably in contact with *R. solanacearum* species (2). Optimal growth conditions for tomato vary between 13 and 33°C depending on the variety studied (37), while the optimum growth temperature for the *R. solanacearum* GMI1000 strain is 28°C (43). Therefore, we hypothesized that panel accessions might have developed defense-related mechanisms specific to their biomes. So, by using a specific tomato wild relative collection of accessions infected at two contrasting temperatures, to mimic temperature elevation under controlled conditions, we aimed to uncover thermostable QDR to *R. solanacearum.* To this end, a panel of 189 tomato wild relatives was selected, sequenced and mapped onto two reference genomes, generated from accessions of two tomato wild relatives representative of the genetic structure of the panel and originating from contrasting climates. Analysis of the population structure of the panel revealed its significant genetic diversity and a subdivision into five clusters. Furthermore, inoculation with the GMI1000 strain at 28°C and 32°C revealed higher response variability and more severe disease at the higher temperature. GWAS conducted following good practices (44), with both reference genomes, at both temperatures, using phenotypic data acquired from a symptom scoring time-course experiment, enabled detection of 28 biologically relevant *QTLs* underlying 44 genes. Our study also highlights the temporal dynamic of plant responses at both temperatures. Most of these *QTLs* were found using both reference genomes - except when genomic structural variations impaired *QTL* detection, however none was common to both temperatures. Interestingly, two *QTLs* are localized near two previously detected major *QTLs*: *bwr-6* and *bwr-12*, for which genes have not been identified.

## Material & methods

### Biological material

#### Tomato panel composition and R. solanacearum strains

Initially, a panel of 199 accessions was constituted, corresponding to a collection of plants available in the Tomato Genetic Ressource Center (University of California, Davis campus, USA) provided by SYNGENTA for the BURNED / Rethink project (INRA n°15000441) and used in phenotyping experiments. Of the original panel, 189 accessions were selected for GWAS. The list of accessions and passport data of the panel are available in **[S1]**. Passport data, when available, include geopositioning (168 accessions), altitude (125 accessions), country of origin, sampling province and species.

The whole panel was inoculated using the wild type *R. solanacearum* GMI1000 strain classified by Hong et al. (45) as a phylotype I, race 1, biovar 3 strain. It originates from South America (French Guiana) (46) and was isolated in tomato (47).

### Plant phenotyping conditions

For each accession, seeds were surface sterilized. Five to 25 seeds were immersed for 10 min in a 15 mL Falcon^TM^ tube filled with 5 mL of a 20 % bleach solution mixed with 200 µL of TWEEN 20. The seeds were then washed four times with the same volume of sterilized water, for 48 h at 4°C for stratification. Seeds were sown on rich BG solid medium (48) in upside down petri dishes for 48 h at 28°C in the dark for germination. A maximum of two germinated seeds per accession were transplanted into pots (7x7x6.4 cm) filled with loam (Proveen potting soil Semi bouturage 2 with perlite, Gergy Fourniture, France). The accessions were grown for two weeks at the Toulouse Plant Microbe Phenotyping plateform (INRAE- LIPME, Toulouse, France) under controlled conditions in a growth chamber [25°C day / 24°C night, 75% RH, 12 h light, LED lights (intensity in μmol m^−2^ s^−1^ per range of wavelengths respectively at 100% Aria: 176; 400-700 / 1; 380-399 / 61; 400-499 / 20; 500-599 / 95; 600-699/ 4; 700-780 / blue, green and peak at 444, 597 and 661 nm)]. Plants were randomized at the beginning of the second week. Five days prior to inoculation, they were moved to a growth chamber in the quarantine laboratory for acclimatization under controlled conditions (28°C day / 27°C night, 75 % RH, 12 h light, 100µmol m^−2^ s^−1^). Inoculations were carried out according to Morel et al. (49), with 40 mL of a solution at 5.10^7^ Colony-forming unit (CFU)/mL/pot. Temperature treatments were applied simultaneously with the *R. solanacearum* inoculation, plants being transferred to growth chambers with the following controlled conditions: 28°C day / 27°C night, 75 % RH, 12 h light, 50 µmol m^−2^ s^−1^ or 32°C day / 31°C night, 75 % RH, 12 h light, 50 µmol m^−2^ s^−1^. The wilting symptoms were scored on an established zero-to-four disease scale as previously described (49) and were monitored from two to ten days after inoculation (Dai).

### Tomato genomic resource creation and analysis

#### Panel sequencing and genome assembly

The panel accessions were sequenced using Illumina chemistry by SYNGENTA (GENEWIZ, US) using Illumina chemistry, with the aim of obtaining a raw sequencing depth between 15X to 25X per accession. The LA1246 and LA1670 accessions, representative of the two genetic groups previously identified in Blanca et al (50), were selected to assemble new reference genomes and to fulfill this purpose were sequenced with a PacBio Sequel Sequencer (Pacific Biosciences, Menlo Park, CA, USA) to reach a minimum depth of 50X. A meta-assembly strategy similar to the one used to assemble the Rosa genome (51) was applied. Very briefly, initial contigs were built using CANU 1.8 (52) and secondary contigs were scaffolded using Bionano optical maps (DLE-1 enzyme, Saphyr system, hybridScaffold.pl software). Details comprising protocols of DNA sample preparation, sequencing and data processing after sequencing are available in **[S2, S3]**.

### Gene prediction & functional annotation

For both reference genomes, protein and non-protein coding gene models were predicted using the integrative EuGene pipeline release 1.5 (53), http://eugene.toulouse.inra.fr/Downloads/egnep-Linux-x86_64.1.5.tar.gz) and five protein databases were aligned with NCBI-BLASTx to detect translated regions. Protein coding genes were annotated by integrating the five sources of information. Results were successively integrated depending on the expected accuracy of the source of information. Priority was given to BLAST-P results from the “Solanum” database **[S2, S3]**.

### Variant calling and selection of Single Nucleotide Polymorphisms (SNP) for downstream analysis

Once genome assembly was complete, variant calling was carried out using FreeBayes (54) on the two genomes that will hereafter be referred to as reference genomes. Several filters were applied to SNP matrices to remove SNPs unsuitable for further analyses **[Table 1]**. Filters A, B, C & D were applied using PLINK1.07 (55). A MAF thresholding of 0.032 was applied according to previously described formula (56). Since preliminary tests showed an over- representation of heterozygous SNPs at extreme GWAS p-values, the filter E was added to avoid disequilibrium between classes of allelic state. It led to the retention of 3,728,946 SNPs (LA1670) and 3,464,481 SNPs (LA1246) for the GWAS SNP matrices **[Table 1].**

**Table 1:** Filters applied to the SNP matrix and the corresponding number of remaining SNPs at each step. Filter A corresponds to FreeBayes output (54) and Filter E was added to avoid a disequilibrium in the representation of each allelic state (56). Identifiers of the filters used and correspondence; A: raw output from genome assembly; B: vcftools filters to remove low quality SNPs (minQ40, biallelic, max missing 0.95); C: minor allele frequency (MAF) at 0.032; D: LD<0.2; E: Threshold used to ensure the presence, for each SNP, of six to seven individuals for each allelic state.

| Filter used | Reference Genome (number of SNP) |  | Analyses |
| --- | --- | --- | --- |
|  | LA1670 | LA1246 |  |
| A | 58 353 015 | 58 606 561 |  |
| A + biallelic | 47 430 644 | 47 486 684 |  |
| A + B | 25 622 125 | 25 929 267 |  |
| A+B+C | 13 541 872 | 14 551 270 | PCA |
| A+B+C+D | 1 836 148 | 2 567 085 | Kinship |
| A+B+C+E | 3 728 946 | 3 646 481 | GWAS SNPs matrix |

### Kinship estimation and population structure analyses

The kinship matrix was computed using the Balding-Nichols method (57) implemented in EMMAX software (Kang et al. 2010), using a filtered matrix (filters A+B+C+D).

Population structure was investigated using Principal Component Analysis (PCA) with R package ***gdsfmt*** (58) and its function *snpgdsPCA().* Results are available in **[S4]**. Additionally, hierarchical clustering was performed on kinship matrices of both reference genomes. Five clusters were chosen for representation. The fixation index (59) between these clusters was then computed with vcftools using the GWAS SNP matrix **[Table 1]** over windows of 10 kbp.

#### Extent of linkage disequilibrium (LD) analysis

The extent of LD (*r*^2^ measure) was investigated in two ways. Firstly, LD half-decay was computed in a three-step procedure: GWAS matrices were sub-sampled (10% of markers for each chromosome), then *r*^2^ between pairs of markers from 1 bp to 1 Mbp in distance were computed; finally the mean LD between pairs of markers within 1 kbp windows was computed. Secondly, to visualize LD patterns along chromosomes, a kinship-corrected LD (60) was computed using a two-step procedure: 1,000 SNPs were firstly sampled from GWAS matrices for each chromosome, then a kinship-corrected *r*^2^ was computed for all SNP pairs on each chromosome using R package ***LDcorSV*** (61).

### Natural variation of QDR in the tomato panel

#### Experimental design

Five blocks of 450 plants were constructed using a random complete block design with two technical repetitions for each block. Each repetition was composed of nine trays of 25 positions corresponding to 225 plants with one plant per accession (*n* = 198 accessions) and the tolerant control accession ‘Hawaii 7996’ (62) placed at the same three positions (top-left corner, center, bottom-right corner) within each tray (*n =* 9 trays x 3 replicates per tray=27).

### Correction for environmental effects and timepoint selection

For each combination of temperature and Dai, referred to as a condition hereafter, phenotypic data were adjusted using the following mixed model (1):

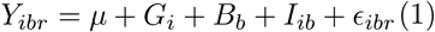

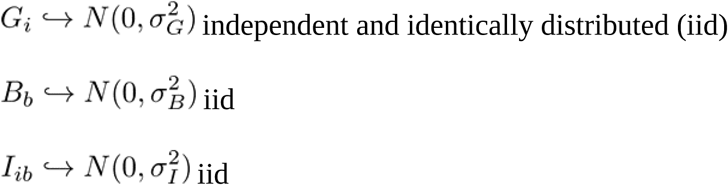

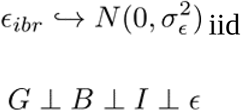

where *Y_ibr_* denotes the phenotype of the *r^th^* repeat of the *i^th^* genotype in the *b^th^* block, *μ* is the intercept, *G* is the genotypic random effect, *B* is the block random effect, *I* is the block x genotype interaction random effect and ε is the vector of errors. ⊥ means independence between random effects. R package ***lme4*** (63) was used to adjust mixed models.

Model (2) was used to compute broad-sense heritabilities, applying the following formula adapted from (64):

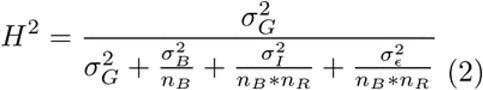

where *n_B_* denotes the number of blocks and *n_R_* the number of repetitions.

Conditions presenting a broad-sense heritability lower than 0.1 were filtered for later analysis **[S5]**. Thus, the following conditions were chosen: 7 and 10 Dai for a temperature of 28°C and 4, 5, 6, 7 and 10 Dai at 32°C. The early days did not pass the threshold since the disease symptoms took a few days to appear.

For the selected conditions and for each accession, least-square-means (lsmeans) were obtained using a modified version of model (1): the genotypic random effect of model was turned into a fixed effect. Lsmeans were then computed using the ***lmerTest*** (65) R package. In the rest of the article, these lsmeans will be designated as the phenotypes.

### GWAS and QTL identification

For each selected condition, the GWAS model from (66) was fitted:

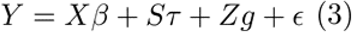

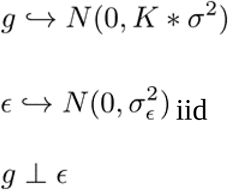

where *Y* is the phenotype vector; τ is the fixed SNP effect; *S* is the SNP incidence matrix (with values in {-1,0,1}); *β* is the fixed structure effect; *X* is the structure matrix (corresponding to the first 3 PCA axes, refer to **[S4]** for the choice of the number of axes); *K* is the kinship matrix; *g* is the random genotypic effect vector and ε corresponds to the vector of errors.

In this study, an approximation for the variance component estimation of the original model from Yu et al. (66) was used for computational purposes, using EMMAX (Efficient Mixed-Model Association eXpedited) (67). We used the EMMAX version implemented in R package ***rrBLUP*** (68), throughout the *GWAS()* function.

Subsequently, the previously described local score method (69) was applied to p- values obtained from the EMMAX model. It aims to control the amount of false positives and to delimit QTL positions precisely. According to Bonhomme et al. (69), the parameter ξ, which controls *QTL* detection stringency, was set to ξ = 3. The procedures returns zones, i.e. intervals of genomic positions in which the association is significant. These significant zones will hereafter be called (putative) *QTLs*. One GWAS was performed for every retained condition. For *QTLs* detected on several consecutive days during kinetics, overlapping <u>QTL</u> positions were merged. Based on the genome annotation of our reference genome, physical positions of genes were calculated. When the position of the gene +/-1000 bp overlapped a *QTL*, the gene was defined as a “strict candidate gene”. An interval of “+/- 1000bp” was added to take into account the promoter regions of genes. The latter definition did not take into account the case where a peak was localized in an intergenic region. The closest genes downstream and upstream of the *QTL* were therefore also considered and were defined as “intergenic candidate genes”.

#### Comparisons of QTLs and candidate genes between two reference genomes

The physical position of genes varied between the two reference genomes. To ascertain the similarity of candidate genes, they were subject to BLAST analysis onto each other’s genomes. Genes were considered similar if they were (1) located on the same chromosome; (2) their BLAST-N p-values were closed to zero; (3) they shared a similar annotation; and (4) they had similar genomic environments (i.e genes were found upstream and downstream).

The relevancy of candidate genes was assessed using the following methodology. First, gene ontologies and identification of putative biological functions in which candidate genes underlying *QTLs* might be involved was investigated. For this purpose, coding sequences of candidate genes were extracted and subject to BLAST analysis (BLAST–N) via the Solgenomics site (https://solgenomics.net/tools/blast/; Tomato Genome cDNA ITAG release 4.0). The same strategy was applied to find orthologs in the *A.thaliana* genome using the BLAST–X tool on the TAIR website (https://www.arabidopsis.org/Blast/), selecting candidates with the highest p-values. Then, the spatiotemporal expression of identified genes was explored using expression browser tools for *A.thaliana* and tomato on the Bio-Analytic Resource for Plant Biology website (eplant tools; https://bar.utoronto.ca). Second, comparison between the two reference genomes was performed to determine whether detected *QTLs* were either common or specific and if structural variations could explain the specificity of the *QTLs*. Gene alignment was performed using annotations of both reference genomes. For each pair of paralogs from the two reference genomes, the following process was applied:

(1) start positions of the two genes were set to zero in a script to align candidate genes at the same absolute position; (2) for visualization of putative structural variation, gene structure (exons, 5’, 3’UTR) was mapped onto each genome for gene pairs using the MultiAlin tool (70) and regions of both reference genomes were aligned with IGV genome browser (71) on a window 13 kbp in length starting at 1 kbp before the candidate gene start codon; (3) finally, positions and p-values of SNPs were added to the custom alignment script to aid in discerning whether the gene coverage of SNPs was equivalent for each reference genome and whether top SNPs were in exons. Moreover by visualizing positions, it was possible to determine if peaks were shared between reference genomes. The *QTL* identified **[Table 3]** were named as follows, using the example of *QTL* ‘9-2-BOTH’: the first number (9) indicates the chromosome, the second (2) indicates the order of appearance on the chromosome (starting from the beginning of the chromosome) and the presence of BOTH indicates that this *QTL* was detected using the two reference genomes. Alternatively, if the *QTL* is specific to one or other of the reference genomes, the pedigree of the latter is indicated. Finally, *QTL* names beginning with ‘IG’ indicate that they have been classified as ‘Intergenic *QTL*’.

**Table 3:** *QTL* and corresponding candidate genes. **A**: name of the *QTL*. **B**: the reference genome(s) on which the *QTL* was detected. According to our candidate gene filters, a * stipulates the *QTL* underlying the candidate gene was detected with one reference genome and was slightly below the threshold p-value with the other reference genome, even if an association peak was observed (see Suppl. Fig **[S11]** for details). **C**: temperature condition (°C) at which the *QTL* was detected. **D**: Corresponding *Solanum lycopersicum* Gene ID and its related annotation found for the ‘Heinz’ reference. **E**: Summary of gene expression pattern in organs retrieved from eplant tomato visualization tools (https://bar.utoronto.ca/eplant_tomato/). **F**: Corresponding *A. thaliana* orthologous gene ID and its related annotation when available; (?) corresponds to genes with low-confidence BLAST results. **G**: Summary of gene expression data in organs retrieved from eplant *A. thaliana* visualization tools (https://bar.utoronto.ca/eplant/). NA: not available.

| QTL name (A) | Reference genome (B) | Temperature (C) | Homolog (D) | Expression pattern (E) | Ortholog (F) | Expression pattern (G) | References |
| --- | --- | --- | --- | --- | --- | --- | --- |
| 1-1-LA1670 | LA1246*/LA1670 | 28 | **Solyc01g006660**<br>Subtilisin-like serine protease, CO(2)-response secreted protease-like | **Roots**, extracellular | **At1g20160**<br>SBT5.2 subtilisin-like serine protease | Stem, roots; **roots**; endoplasmic reticulum, extracellular; **negative response to bacteria** | (Ascencio-Ibanez et al., 2008) |
| 1-2-LA1246 | LA1246/LA1670* | 32 | **Solyc01g086890**<br>Splicing factor 4-like protein G-patch | Leaves, mature fruits, **roots**; nucleus | **At3g52120**<br>CIS1, CRY2 interacting splicing factor 1, regulate thermosensory flowering via FLM alternative splicing. | Everywhere; nucleus, cytosol |  |
| 1-3-LA1246 | LA1246/LA1670* | 32 | **Solyc01g100630**<br>Catalase | Leaves, fruits, **roots**, peroxysome | **At4g35090 (?)**<br>multigene family | everywhere; peroxysome, mitochondrion, **response to infection** |  |
| 1-3-LA1246 | LA1246/LA1670* | 32 | **Solyc01g100640**<br>Catalase | fruits, peroxysome | **At4g35090 (?)**<br>multigene family | everywhere; peroxysome, mitochondrion, **response to infection** |  |
| 1-3-LA1246 | LA1246/LA1670* | 32 | **Solyc01g100650 (?)**<br>NHL repeat-containing protein 2-like | leaves, fruits, chloroplast | **At1g56500** | everywhere; plastids; small response to pathogen infection |  |
| 1-4-BOTH | LA1246/LA1670 | 28 | **Solyc01g105260**<br>Protein of unknown function DUF125, vacuolar iron transporter homolog 4-like | **Roots** | NA | NA |  |
| 1-4-BOTH | LA1246/LA1670 | 28 | **Solyc01g105270**<br>SITOM1b tobamovirus multiplication 1 homolog b | Mature fruits and **roots**; plasma membrane | NA | NA | (Ishikawa et al. 2022) |
| 2-1-BOTH | LA1246/LA1670 | 28 | **Solyc02g067460**<br>Mitochondrial outer membrane protein porin | **seedling roots**, fruits, mitochondrion | Multigene family | NA |  |
| 3-1-BOTH | LA1246/LA1670 | 32 | **Solyc03g123790**<br>Putative calcium/proton exchanger | **roots**, fruits, plasma membrane | **At1g55730 (?)**<br>multigene family | NA |  |
| 3-2-LA1670 | LA1246*/LA1670 | 28 | **Solyc03g093130 (?)**<br>xyloglucan-modifying enzyme | Fruits, **roots**; extracellular | **Multigene family** | NA |  |
| 3-3-LA1670 | LA1246*/LA1670 | 32 | **Solyc03g097470**<br>3-Beta-ketoacyl-acyl carrier protein synthase III (FabH) | Fruits; plastids | **At1g62640**<br>3-ketoacyl-acyl carrier protein synthase III | Everywhere; **response to infections** | (Ascencio-Ibanez et al., 2008) |
| 3-4-LA1670 | LA1246*/LA1670 | 28 | **Solyc03g019730**<br>Sumo activating enzyme 1b Molybdenum cofactor biosynthesis | Leaves, fruits, **roots**; mitochondria | **At5g50580 & At5g50680**<br>AT-*SAE1-2*, *SAE1B*, sumo-activating enzyme 1B | Seeds; cytosol |  |
| 4-1-LA1246 | LA1246/LA1670* | 28 | **Solyc04g008310**<br>UDP-glucuronosyl/UDP-glucosyltransferase | Leaves, fruits, **roots** | **At2g15490**<br>UDP-glycosyltransferase 73B4 | Seeds, **roots**; cytosol, cell wall, **response to biotic and abiotic stresses** | (Langlois-Meurinne et al., 2005) |
| 4-2-BOTH | LA1246/LA1670 | 28 | **Solyc04g078590**<br>Receptor kinase Serine/threonine kinase | Flowers, immature fruits; plasma membrane | **Multigene family** | NA | [Gou et al., 2023] |
| 7-1-LA1670 | LA1670 | 32 | **Solyc07g005040 & Solyc00g257110**<br>ATPase 8, plasma membrane-type-like | Flowers | **Multigene family** | NA |  |
| 7-2-BOTH | LA1246/LA1670 | 32 | **NA** | NA | **NA** | NA |  |
| 7-3-LA1670 | LA1246*/LA1670 | 28 | **Solyc07g044730 (?)**<br>3-hydroxyisobutyryl-CoA hydrolase-like protein 5 | flowers (?) | **At1g06550 (?)**<br>multigene family | NA |  |
| 8-1-BOTH | LA1246/LA1670 | 32 | **NA** | NA | **NA** | NA |  |
| 9-1-BOTH | LA1246/LA1670 | 32 | **Solyc09g009700**<br>GDSL esterase/lipase | Flowers, **roots**; extracellular | **At5g03610**<br>GGL25 - GDSL-motif esterase/acyltransferase/lipase | Stem, embryo, **primary root**; endoplasmic reticulum, extracellular; **response to pathogen** and nitrogen | (Mohr & Cahill, 2007) |
| 9-2-BOTH | LA1246/LA1670 | 32 | **Solyc09g011640**<br>Glutathione S-transferase-like | Leaves, **roots**; cytosol | **At2g29420 (?)**<br>Glutathione S-transferaseE | **Roots**, petals. **Response to hormones and infection** |  |
| 10-1-LA1246 | LA1246/LA1670* | 32 | **Solyc10g081260**<br>Multidrug resistance protein mdtK/ Multi antimicrobial extrusion protein MatE | Fruits; plasma membrane | **At5g17700**<br>MATE efflux family protein | **Roots**, petals; plasma membrane; strong **response to infections** | (Ascencio-Ibanez et al., 2008) |
| IG1-1-LA1670 | LA1246*/LA1670 | 28 | **Solyc01g016350**<br>(Unknown Protein) | NA | NA | NA |  |
| IG1-1-LA1670 | LA1246*/LA1670 | 28 | **NA** | NA | NA | NA |  |
| IG1-1-LA1246 | LA1246/LA1670* | 28 | **NA** | NA | NA | NA |  |
| IG1-1-LA1246 | LA1246/LA1670* | 28 | **NA** | NA | NA | NA |  |
| IG2-1-LA1670 | LA1670 | 28 | **Solyc02g084680**<br>Protein SHI RELATED SEQUENCE 5 | flower buds, fruits, **roots** | multigene family | NA |  |
| IG2-1-LA1670 | LA1670 | 28 | **Solyc02g084670**<br>NA | flower buds, plastids | multigene family | NA |  |
| IG3-1-LA1246 | LA1246 | 28 | **Solyc03g093770**<br>histone-lysine N-methyltransferase | leaves ? | NA | NA |  |
| IG3-1-LA1246 | LA1246 | 28 | **NA** | NA | **At4g12560 (?)** | NA |  |
| IG4-1-LA1246 | LA1246/LA1670* | 32 | **Multigene family** | NA | **Multigene family** | NA |  |
| IG4-1-LA1246 | LA1246/LA1670* | 32 | **Solyc00g028960**<br>Glutaredoxin family protein | leaves | multigene family | NA |  |
| IG4-1-LA1246 | LA1246/LA1670* | 32 | **NA** | NA | **NA** | NA |  |
| IG6-1-BOTH | LA1246/LA1670 | 32 | **Solyc06g051510**<br>Protein kinase superfamily protein | **roots**, fruits, leaves, nucleus | multigene family | NA |  |
| IG6-1-BOTH | LA1246/LA1670 | 32 | **Multigene family** | NA | **Multigene family** | NA |  |
| IG6-2-BOTH | LA1246/LA1670 | 32 | **Solyc06g065070**<br>FAD-binding domain-containing protein | fruits ?, vacuole | multigene family | NA |  |
| IG6-2-BOTH | LA1246/LA1670 | 32 | **Solyc06g065060**<br>FAD-binding domain-containing protein | fruits ?, vacuole | multigene family | NA |  |
| IG7-1-LA1670 | LA1246*/LA1670 | 32 | **Solyc07g007610**<br>At2G21500 protein (Fragment) (AHRD V1 *--- B9DHX0_ARATH); contains Interpro domain(s) IPR001841 Zinc finger, RING-type | flower buds, nucleus | multigene family | NA |  |
| IG7-1-LA1670 | LA1246*/LA1670 | 32 | **Solyc07g007600**<br>vacuolar-type H+-pyrophosphatase | fruits, **roots**, leaves | **At1g15690** | Everywhere; mostly vacuole & plasma membrane; response to some infections |  |
| IG7-2-BOTH | LA1246/LA1670 | 32 | **NA** | NA | **NA** | NA |  |
| IG7-2-BOTH | LA1246/LA1670 | 32 | **Solyc07g007940**<br>Small nuclear RNA activating complex (SNAPc), subunit SNAP43 protein (AHRD V3.3 *** AT3G53270.9) | fruits, **roots**, leaves, cytosol | **At3g53270** | Everywhere; cytosol, nucleus; response to some infections | (Ascencio-Ibanez et al., 2008) |
| IG8-1-LA1670 | LA1670 | 32 | **Solyc08g045620**<br>Ycf2 (AHRD V1 *--- A6Y9Y9_9MAGN); contains Interpro domain(s) IPR008543 Chloroplast Ycf2 | NA | **AtCg01280 & AtCg00860** | Everywhere; mostly plastids; small response to infections |  |
| IG8-1-LA1670 | LA1670 | 32 | **NA** | NA | **NA** | NA |  |
| IG12-1-BOTH | LA1246/LA1670 | 32 | **Solyc12g008960**<br>Calmodulin-binding protein | **roots**, nucleus | **At4g33050 (?)**<br>multigene family | Everywhere; cytosol, mitochondrion, nucleus; **response to some infections** |  |
| IG12-1-BOTH | LA1246/LA1670 | 32 | **Solyc12g008950**<br>DEAD-box ATP-dependent RNA helicase | **roots**, nucleus | **At1g72690 (?)** | nucleus; response to some infections |  |

## Results

### Panel characteristics and choice of reference genomes

The 189-accession panel used in this study included five *Solanum* species: 167 *S. pimpinellifolium*, 20 *S. lycopersicum* var *cerasiforme,* one *S. cheesmaniae*, and the *S. lycopersicum* cultivar ‘Hawaii7996’, known to confer tolerance to bacterial wilt (62). The precise localization information available for 168 accessions enabled their positioning on a map **[Fig. 1, A and C]**. Among the 189 accessions selected, a majority came from Peru (73%) and a significant proportion came from Ecuador (14 %). The *S. lycopersicum* var. *cerasiforme* accessions originated from Central America and from the western side of the Andes. Some *S. pimpinellifolium* were also found in this area, but most were sampled from the eastern side, either in the mountainous areas of southern Ecuador / northern Peru or in the coastal valley near the Pacific Ocean in southern Peru. The accessions were collected at different altitudes. The majority (45 %) were found below 500 m. 11 % were found in a range of altitudes between 500 m and 1,000 m and finally 11 % above 1,000 m. The highest sampling point indicated was 3,000 m.

**Figure 1:**
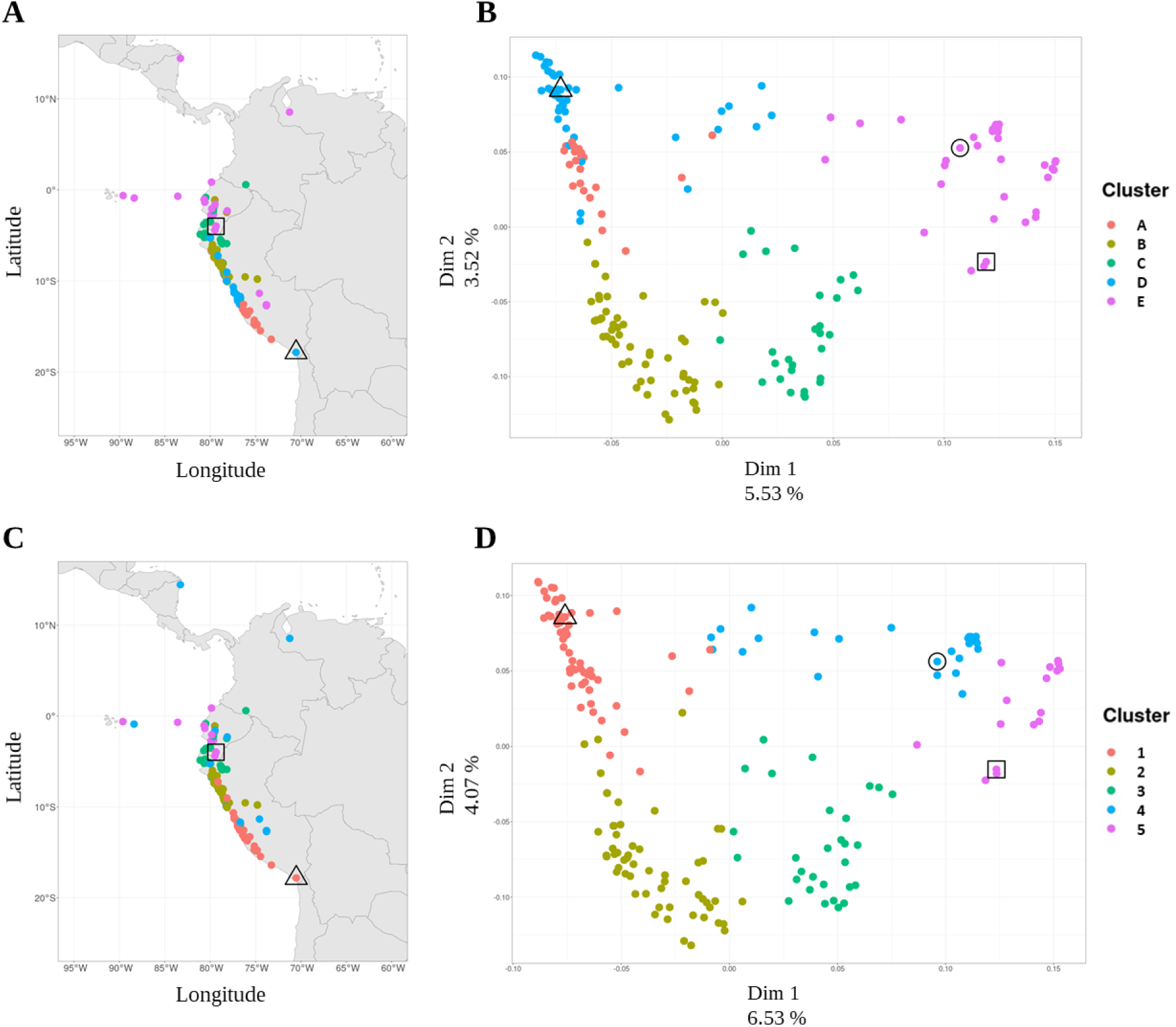
Geographical origin and population structure of the panel. **A**-**C**: Geographical localization on a map of South America of the groups identified depending on the reference genome used: LA1246 (**A**) or LA1670 (**C**). **B**-**D**: Representation of the panel in the first two PCA axes computed with population structure using LA1246 (**B**) and LA1670 (**D**) reference genomes. Colored spots represent groups of accessions that were identified using kinship clustering (see the Material & methods section for more details). Square: LA1246 from Ecuador; triangle: LA1670 from Peru; circle: cultivar ‘Hawaii 7996’.

We selected two reference accessions within our panel based on results from a previous genetic study and climatic data obtained using their GPS coordinates (72), [**S1, S6].** Indeed, Blanca et al. showed that genetic diversity of their smaller panel of *S. pimpinellifolium* was organised in two major genetic clusters (50). We decided to select one accession per cluster and in addition accessions had to belong to two contrasted biomes. Thus, we selected LA1670 and LA1246: LA1670 originated from an arid environment and LA1246 a wetter cooler environment.

### Genomic data analyses of the panel

Reference genomes were sequenced using PacBio technology and reached a depth of 82X for LA1246 and 80X for LA1670 **[S2, S3]**. The results obtained were similar for both reference genomes and were evidence of their high sequence quality. Interestingly, genome assembly highlighted large chromosomic rearrangements between the two reference genomes, especially on chromosomes 6, 7 and 8. The final assembled genome size was 815.7 Mb for LA1670 and 808.6Mb for LA1246. Gene prediction revealed 46,202 (LA1670) and 45,746 (LA1246) genes.

Illumina sequencing of the panel revealed a depth of between 15X and 25X for raw sequenced data which varied among accessions **[S2]**. For SNP **[Table 1, filter A]**, the average coverage at a locus was between 13.66x (LA1246) and 13.81x (LA1670).

Reference genomes were used to call SNPs obtained by panel sequencing. We assessed the gene SNP coverage by using filtered GWAS SNP matrices **[Table 1, filter A, B, C and E]**. Results indicated that at least one SNP was found in 80% of genes with, on average, 9.6 SNPs/gene whichever reference genome was chosen. This coverage varied between 7.5 and 13.2 SNPs/gene depending on the chromosome **[Fig 2]**.

**Figure 2:**
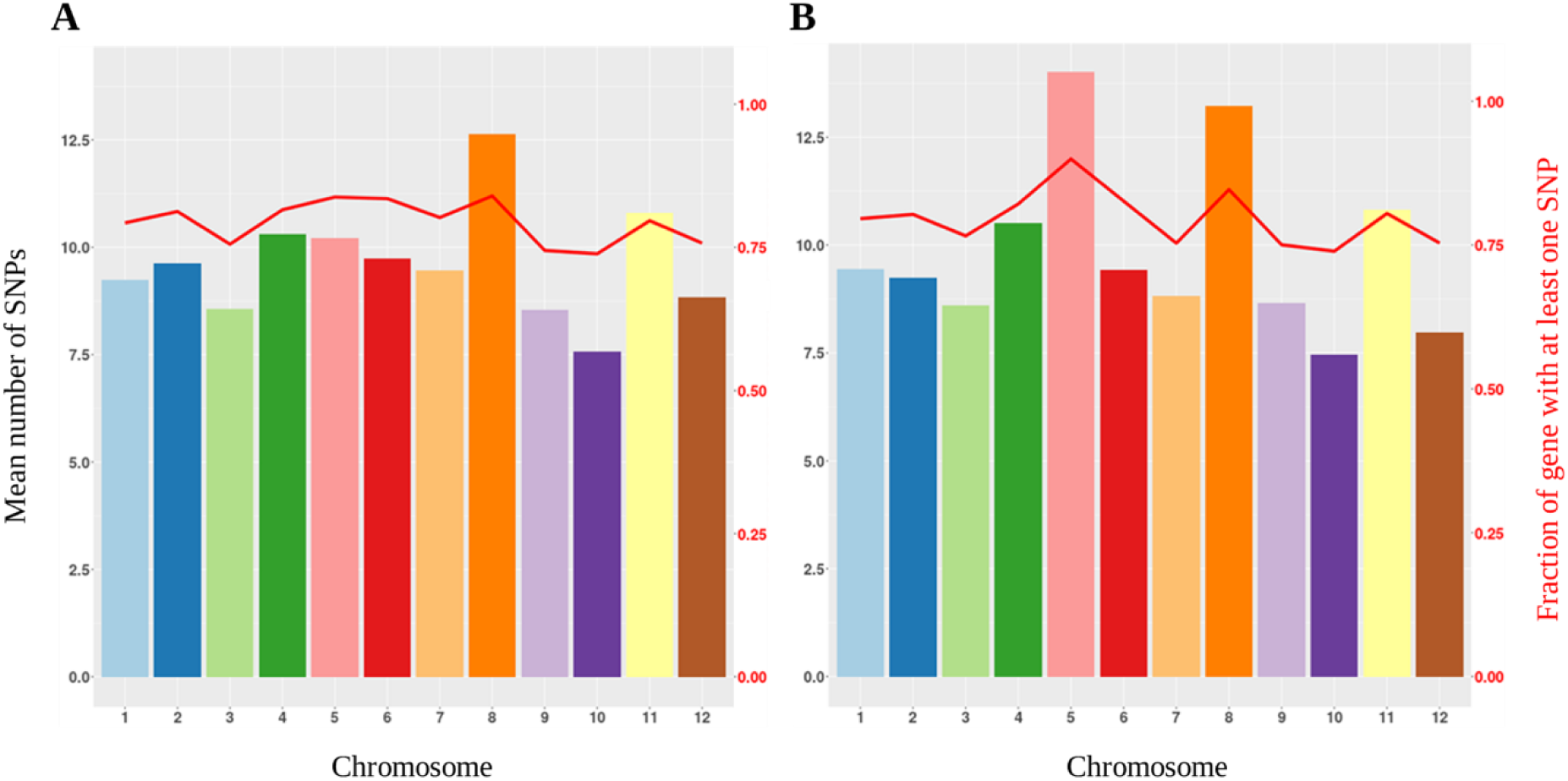
Gene coverage by chromosome for each reference genome. Bar height indicates the mean number of SNPs per gene for each chromosome. **A:** LA1246, **B:** LA1670. The red line indicates the fraction of the gene containing at least one SNP.

### Population structure and LD analysis reveals a large panel genetic diversity suitable for GWAS

Population structure was investigated through PCA using SNPs called from the two reference genomes. **[Fig. 1, B and D]** represent panel accessions on the first two PCA axes, which explain cumulatively around 10% (9.05% for LA1246 and 10.60% for LA1670) of the genetic variation, along which panel accessions segregate into five clusters. These results indicate that the choice of the reference genome has little impact on the PCA and on the clustering. Based on their distance on PCA axes **[Fig. 1, B and D]** the two reference genomes appear diversified.

The first PCA axis can be interpreted as a North-South gradient whereas the second PCA axis suggests the existence of an East-West gradient **[Fig. 1, A and C]**. Accessions in the middle of the graph **[Fig. 1, B and D]** are probably admixed genotypes. Clusters identified from the hierarchical kinship clustering are represented in colour **[Fig. 1]**, and show consistent F ST values **[S7]**. For instance, accessions from clusters 4 and 5 for LA1670 are close together on the PCA axes **[Fig. 1D]**, on a geographical map **[Fig. 1C],** and share a low F ST value **[S7]**. The extent of LD was investigated using the filtered GWAS matrix **[Table 1, filters A, B, C and E]**; its half-decay was calculated per chromosome and is of order of a few kbp for most chromosomes, with the exception of chromosomes 5, 6 and 8 for which LD decay is lower **[S8, S9]**. Specifically, LD visualization on these latter chromosomes showed large blocks of LD, located in central chromosomic regions, whereas LD is relatively low in telomeric regions **[S10]**.

### Panel responses to R. solanacearum reveal three temperature-dependent profiles

Panel accessions were phenotyped at 28°C and 32°C for the severity of bacterial wilt disease symptoms for ten days following pathogen inoculation. The three temperature- dependent profiles identified among panel accessions and shown in **[Fig. 3]** were: (1) accessions more susceptible to disease at 32°C than at 28°C (133 out of 189 accessions); (2) accessions exhibiting a same behavior independently of the temperature applied (49 out of 189); (3) accessions less susceptible at 32°C (7 out of 189) **[Fig. 3]**. Following inoculation, disease severity increased with time **[Fig. 4]**, indicating that plants became gradually more susceptible and could not recover from the disease. At 28°C, the trend showed a difference in response to *R. solanacearum* between the two reference accessions, with LA1246 being symptomless **[Fig. 4]**. However, at 32°C, both reference accessions developed wilting symptoms. The ‘Hawaii 7996’ cultivar, which was used as a control due to its tolerance to bacterial wilt, started to wilt moderately in the later days of the experiment at 32°C **[Fig. 4]**. Interestingly, for some accessions, disease symptom curves remained below the one of ‘Hawaii 7996’ **[Fig. 3]**, suggesting that these accessions were more resistant to bacterial wilt than ‘Hawaii 7996’ at 32°C. Altogether, phenotypic data suggested temperature-dependent patterns of response to *R. solanacearum*.

**Figure 3:**
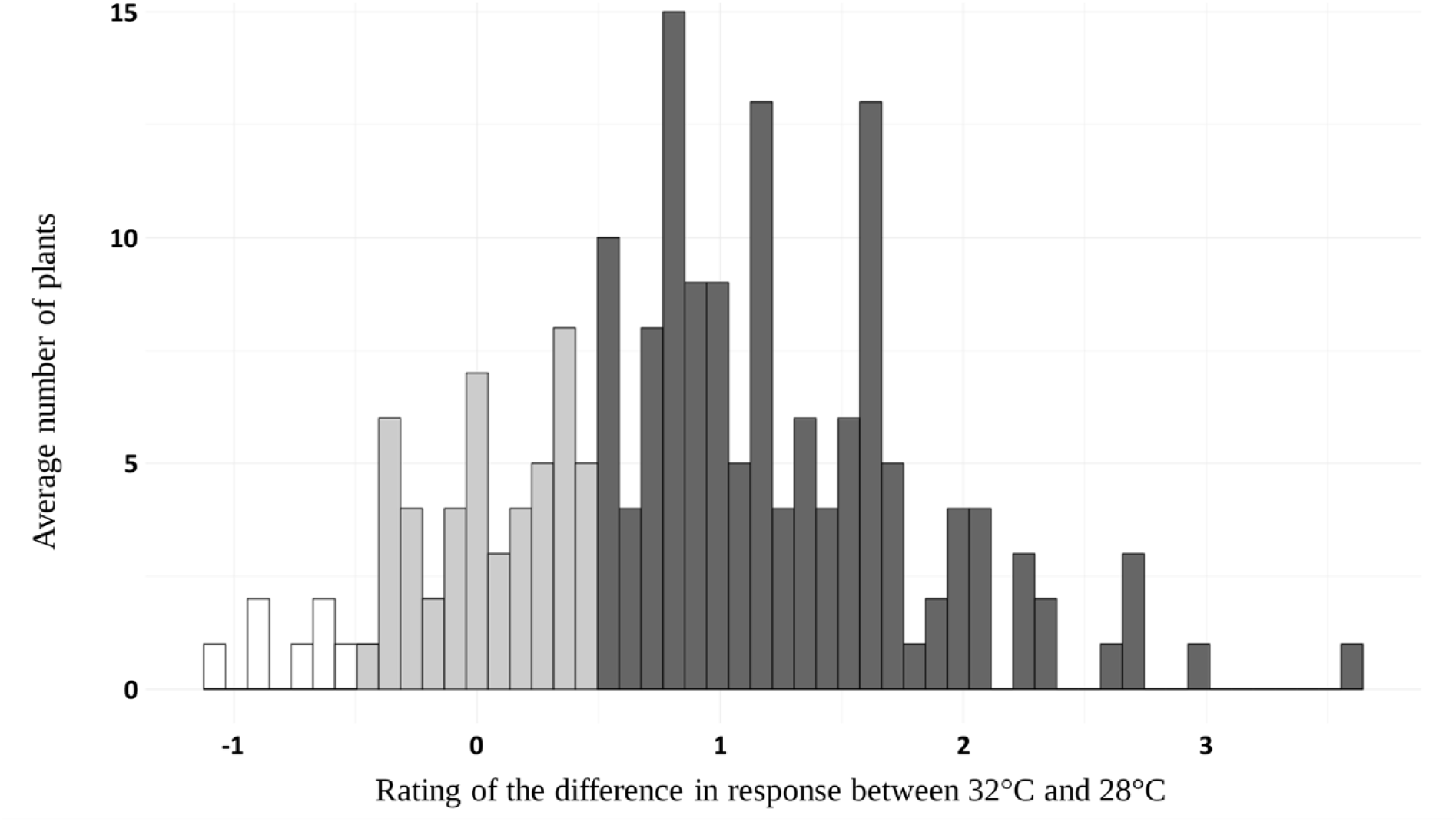
Difference between the phenotypic response of the panel to *R. solanacearum* at 32°C and 28°C at 10 Dai. Ratings used were adjusted means (lsmeans) for every panel accession, which enabled classification of the panel into three groups according to their defense response to *R. solanacearum*. Colours represent groups in which the response to *R. solanacearum* differs. Dark gray: more susceptible at 32°C 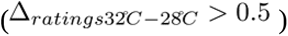; light gray: similar at both temperatures 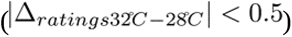; white: more susceptible at 28°C (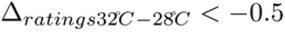).

**Figure 4:**
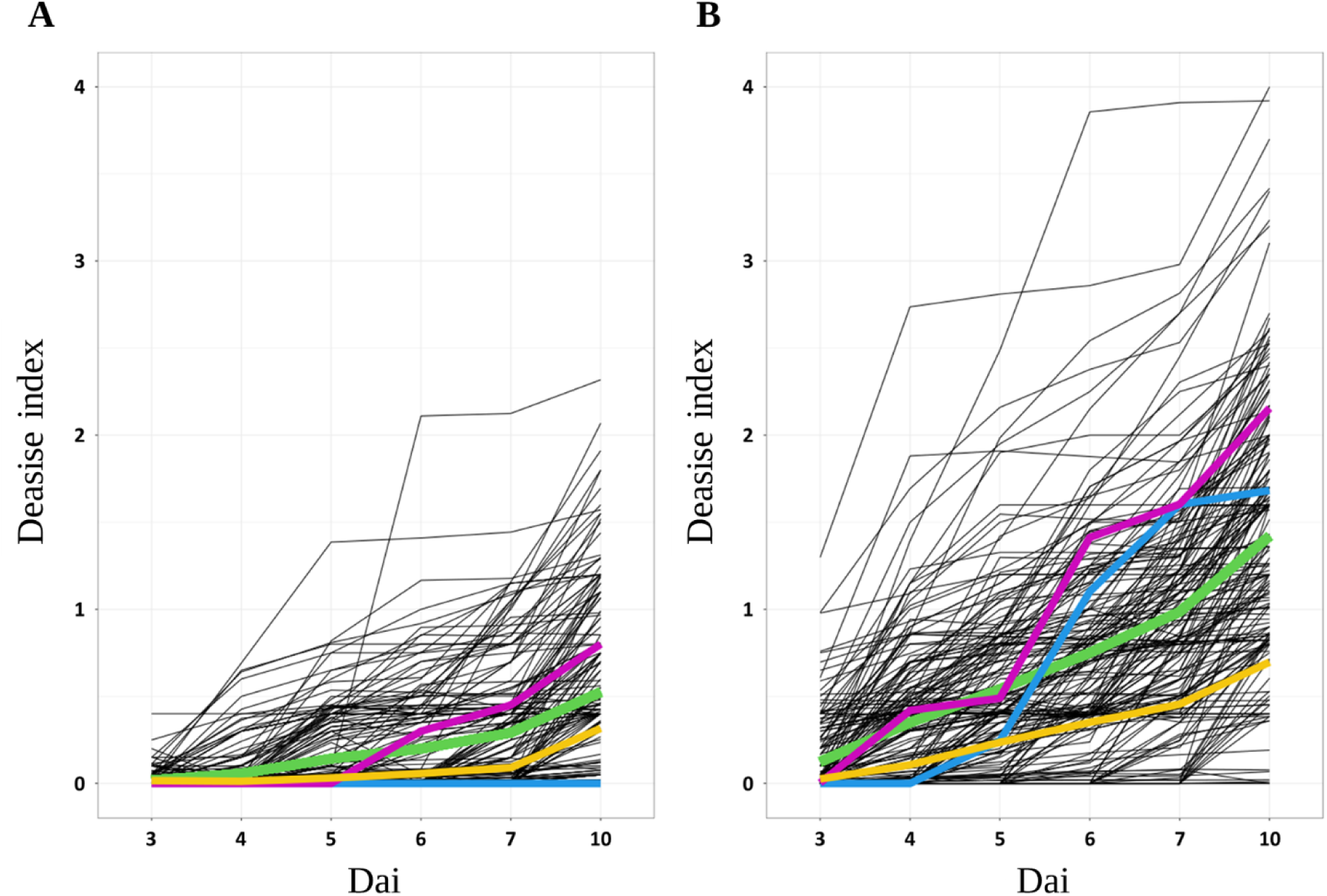
Kinetics of the response of panel accessions to *R. solanacearum* at two different temperatures. **A**: 28°C; **B**: 32°C. The y-axis represents adjusted means (Lsmeans) of the disease index (ranging from zero to four) over Dai. Each line represents the disease curve of one accession. Coloured curves represent the overall mean response of the panel (green), the ‘Hawaii 7996’ cultivar (yellow) and the accessions used as reference, LA1246 (blue) and LA1670 (purple).

### GWAS and post-GWAS analyses identify relevant candidate genes involved in bacterial wilt defense response

GWA was performed on a sub-sample of selected conditions (see M&M). Results are presented in **[S11]** and **[Table 3]**. After filtering *QTLs*, 21 “strict candidate genes” (see M&M for details) belonging to 18 *QTLs* were selected. *QTL* size did not exceed a few kbp. In almost all cases, one candidate gene was positioned at one *QTL*, indicating gene-level resolution. Exceptions include *QTL 1-3-LA1246* whose position was larger (12 kbp); gene density was particularly important in this region. The other exception was *QTL 1-4-BOTH*, which underlies two candidate genes; however, SNPs with the lowest p-values (RIZ2870_SpimpiLA1670Chr01_88223852_T for LA1670 and RIZ2870_SpimpiLA1246Chr01_89956087_T for LA1246) were located in an exon of the *Solyc01g105270* homolog. When “intergenic candidate genes” were considered **[Table 3]**, 23 additional candidate genes related to ten *QTLs* were added. Even if they did not pass the ‘vicinity with annotated genes’ criterion, some were of interest: for instance, *QTL IG12-1- BOTH* was systematically detected over four consecutive days **[S11]**, which constitutes the maximum number observed in the analysis.

### GWAS using two reference genomes improve QTL detection

Among the 21 “strict candidate genes” presented in **[Table 3]**, 11 out of 21 (indicated by a * in **[Table 3])** were identified with one reference genome and displayed an association peak slightly below the detection threshold with the other one, and 9 were identified with significant peaks in both reference genomes. These 20 candidate genes can be considered as common to the two reference genomes.

First, a statistical explanation could be given: based on the criterion of *QTL* selection, some *QTLs* were close to the significance threshold and passed it only with one reference genome. This is the case for *QTL* 1-2-LA1246, identified as specific to LA1246 **[Fig. 5A]**. We selected this *QTL* because it was detected at only one Dai at 32°C, but its peak exceeded ten in local score (LA1246). By contrast, at the same position on the LA1670 reference genome, the peak value was below ten, also at the same Dai, meaning this *QTL* did not meet the criteria for our selection. However, since the peak surpasses the significance threshold of the local score method (red and blue lines), *QTL 1-2-LA1246* has been flagged as nearly shared (denoted by an *) in **[Table 3]**. The GWAS model used was based on a statistical test using both kinship and population structure (see M&M for more details). Although the correlations between kinship matrices from the two reference genomes were strong (> 0.9), the differences between them could still result in the detection of different *QTLs* in each reference genome.

**Figure 5:**
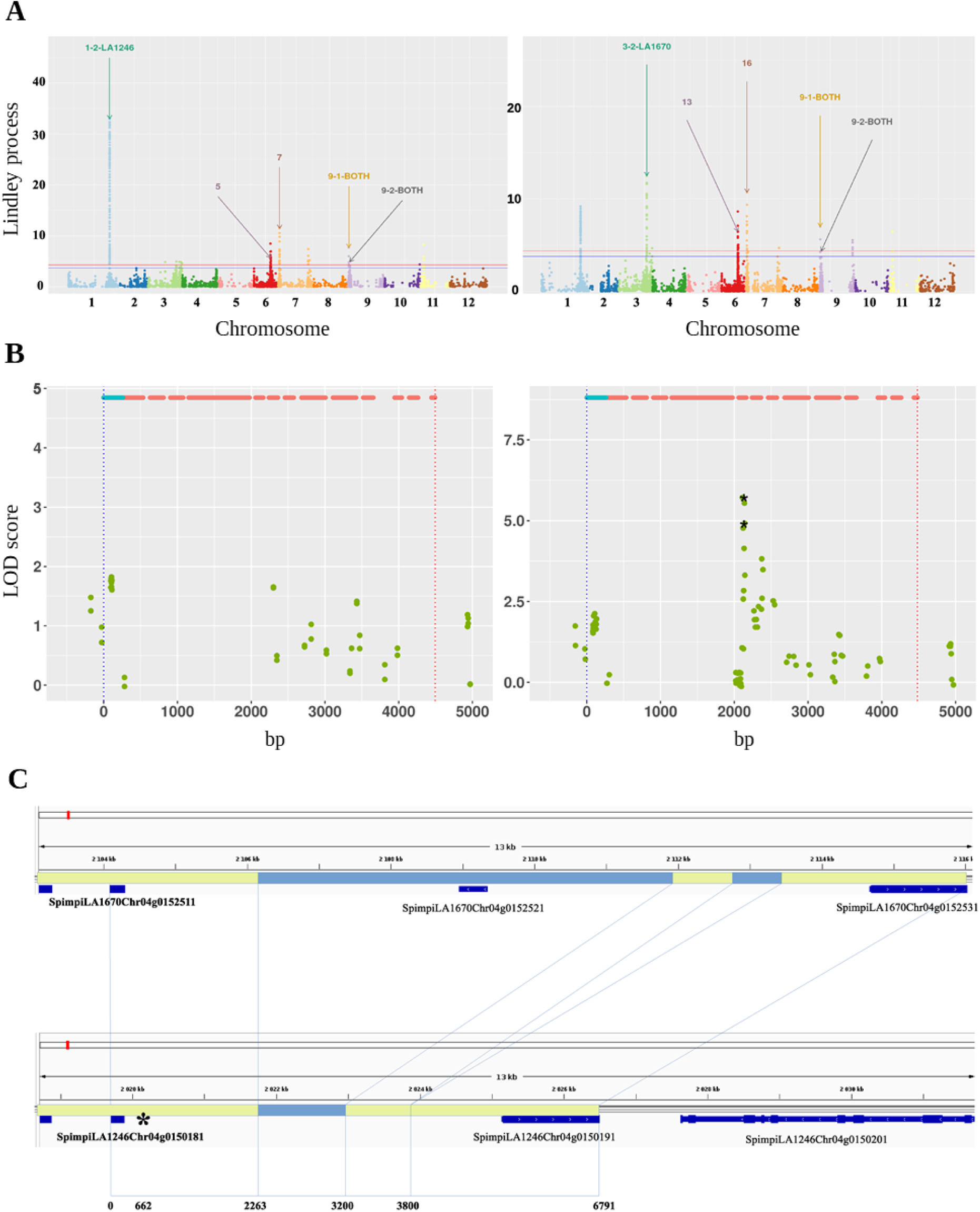
Three different cases for which the use of two reference genomes helped *QTL* detection. **A:** example of the effect of the reference genome on *QTL* detection due to selection criteria. GWAS with local score at 32°C at 10 Dai for LA1246 (left) and LA1670 (right) reference genomes. The light-blue peak in chromosome 1 underlies *QTL 1-2-LA1246*, reported as being unique to LA1246 even if a peak of association, below the threshold p-value of detection, was observed in LA1670 (* in **[Table 3]**). The x-axis represents markers’ chromosomic positions. The y-axis represents local score values. Maximal (red line) and minimal (blue line) detection thresholds for *QTLs* with the local score approach are shown. **B:** example of the *QTL 7-1-LA1670* for which variant calling based on reference genome LA1246 (left panel) failed to detect an SNP in the exon of a *QTL* that was detected using the LA1670 reference genome. SNPs (dots) are localized on the gene sequence whose structure (exon in red, 3’UTR in blue) is shown at the top of each graph. Arrows indicate the exon on which the *QTL* is detected for LA1670. The y-axis represents -log_10_ (p-values). Random noise was added to the x-axis for representation purposes for all p-values in the two conditions. *: The most significant SNP. **C:** example of structural rearrangements between the LA1670 (upper panel) and LA1246 (lower panel) reference genomes with two indels (correspondence of the genomic regions specified in dotted lines) that may have affected detection of the *QTL 4-1-LA1246* counterparts in LA1670. Alignments of common genomic regions with good accuracy are colored in light green while different genomic regions are in light blue.

A second explanation is linked to the variant calling used when building SNP matrices. A candidate gene cannot be detected if no informative allelic variant is present in proximity. [**Fig. 5 B]** illustrates this situation with the *QTL 7-1-LA1670*, underlying a gene encoding an APT8 Plasma membrane like protein with different SNP coverage depending on the reference genome used. The high gene coverage with LA1670 allowed its detection whereas the exon in which the top SNP is found is not covered by any SNP in the LA1246 reference.

A final explanation relates to structural variations between reference genomes, which were investigated by BLAST analysis of reference genome sequences against each other. For instance, structural variations affecting the *QTL* detection are highlighted in **[Fig. 5 C]** for the *QTL 4-1-LA1246* underlying a gene encoding a potential UDP-glycosyltransferase. The two indels modifying the genomic environment on LA1246 reference genome may have affected the detection of its counterpart in LA1670. The differential detection of several *QTLs* specific to one of the two reference genomes, as shown in **[Fig. 5 A]**, could be attributed to this type of structural variation. For example, a genomic inversion is observed only near *QTL 3-2- LA1670*. Structural variations also particularly affected the regions of ’intergenic candidate genes’; for instance, an inversion was detected near *QTL* IG3-1-LA1246, while an indel of approximately 200 kbp was found between the two genes underlying *QTL* I*G8-1-LA1670*. Additionally, BLAST analysis of the genes closest to *QTL IG2-1-LA1670* in the LA1246 genome did not reveal a gene annotated as a putative serine/threonine phosphatase, which is present in the LA1670 genome.

### GWAS highlight temperature and time-dependent, biologically relevant candidate genes

#### Different QTLs detect at 28°C and 32°C

Comparison of GWAS results showed that no *QTL* was common to the two temperatures. Among the 21 ’strict candidate genes’ (see Materials & Methods section for details) corresponding to 18 *QTLs*, a similar number of *QTLs* was identified at both temperatures: eight out of 18 at 28°C and ten out of 18 at 32°C. This suggests that the plant response to GMI1000 is quantitative but regulated differently at the two temperatures.

### Detected QTLs vary over time following pathogen inoculation

Regardless of temperature, peaks appeared and disappeared on Manhattan plots depending on the phenotyping day **[S11]**, suggesting that plants may use different strategies as the infection progresses. Notably, when a *QTL* is present, it is detected on either one or, more commonly, two consecutive days. Exceptions include *QTLs 7-2-BOTH* and I*G8-1- LA1670*, observed over three days, and *QTL IG12-1-BOTH*, detected over four days.

### Biological relevancy of candidate genes underlying detected QTLs

BLAST analyses were performed to find tomato homologs and *A.thaliana* orthologs of candidate genes. Expression patterns were investigated when statistically relevant homologs/orthologs were found by BLAST and when transcriptomic data were available. We focus on “strict candidate genes” because they had a higher GWAS resolution enabling a better identification of the causal gene. Twelve out of 19 tomato homologs were expressed in roots in various cell compartments. Interestingly five genes (*Solyc01g006660*, *Solyc01g105270, Solyc04g008310, Solyc09g009700* and *Solyc10g081260)* have already been annotated as immunity-related genes and involved in response to other pathogens. Only eight orthologs of *A.thaliana* were found with high BLAST confidence levels **[Table 3]**, probably due to the genetic distance between *S. pimpinellifolium* and *A. thaliana*. Five out of eight were particularly expressed in roots and six out of eight showed different expression levels when plants were subjected to a biotic stress. Although many candidate genes could not be analyzed further due to a lack of available data in literature, the trend is towards candidate genes that appear to be preferentially expressed in roots and on pathogen attack.

## Discussion

To identify resistance to *R. solanacearum* that remains efficient at elevated temperature in tomato, we chose to study the pathosystem tomato/*R. solanacearum* at two contrasted temperatures: the one commonly used in controlled conditions (28°C) and also 32°C – this last temperature being chosen to mimic temperature elevation, and having been described as altering the defense response of tolerant tomato cultivars in the field (39). To this end, we implemented a GWAS approach using a selected panel of wild tomato parents composed mainly of *S. pimpinellifolium* accessions, newly sequenced on this occasion, and including two internal reference genomes.

The selection of the panel was based on the fact that tomato ancestors originated from a broad geographical range, spanning Central to South America, and likely evolved and adapted to diverse environments. Indeed, the Central American regions experience an equatorial climate, with elevated average temperatures and abundant rainfall, while the climate in the southern regions is more arid, with colder temperatures and lower precipitation (73). The presence of RSSC in this vast geographical region has been demonstrated by Wicker and colleagues, with phylotype II strains being specifically detected in Central and South America (2). As the main abiotic determinants of bacterial wilt development are temperature and moisture (38), and tomato defense response to bacterial wilt is known to be thermo-sensitive (32, 39), we hypothesized that tomato wild-relatives developed biome- and temperature- dependent defense mechanisms to RSSC. Our phenotyping results confirmed this assumption, with the accessions showing wide variability in their response to the *R. solanacearum* GMI 1000 reference strain, which was further accentuated by elevated temperature **[Fig. 3]**. This finding contrasts with previous results obtained in *A. thaliana,* showing little variation of plant response to the bacteria at 30°C, with nearly all accessions being susceptible ten days after inoculation. They also suggest tomato wild relatives are better adapted to elevated temperatures (20, 22). Interestingly, the evolution of the response between 28°C and 32°C allowed the clustering of the accessions into three distinct groups, with a majority of accessions becoming more susceptible at the elevated temperature. Another fraction is more resistant, and the final group shows the same response independently of the temperature applied. These results support the existence of diverse underlying genetic architectures of plant responses to combined stress within the panel **[Fig. 4]**.

The analysis of population structure **[Fig. 1]** confirmed the wide genetic diversity of the panel, raising the question of the choice of an adequate reference genome for variant calling. In tomato, the majority of GWAS papers have used the ‘Heinz’ cultivar as the reference genome (74, 75). However, ‘Heinz’ is a cultivar that has experienced a reduction in genetic diversity due to domestication (76). Indeed, immunity and plant development is a trade-off that plants have to face with a limited quantity of energy. Some parts of the genome, notably immunity genes, may have suffered a reduction compared to wild relatives (variant copy number, indels, absence of genes etc.) (76). Consequently, when carrying out variant calling, some genomic regions, potentially linked with immunity are either poorly or not covered by SNPs. A natural alternative to the Heinz genome would be to use a pangenome as a reference for variant calling. Indeed, pangenomes aim to capture as much structural variation as possible within a species, making them an optimal choice for consistent variant calling (77). However, constructing a pangenome requires large amounts of high-quality data and significant bioinformatics resources (78). In this article, we propose an alternative strategy that involves assembling two reference genomes selected within the panel, an approach that remains relatively uncommon in GWAS (79).

Our proposal is first to select reference genomes that are relevant to the biological question. In the present study the objective was to identify mechanisms that remain effective in fluctuating environments, especially in the case of temperature elevation. Consequently, the two reference genomes selected originated from contrasted climates **[S6].** The choice was also influenced by their geographical and genetic distance **[Fig. 2]**. As with a pangenome, the aim is to be able to call variants in all regions of the two reference genomes, and more particularly to be able to cover non-homologous regions. For GWAS, the reference genome chosen affects GWAS outputs by modifying peak height on Manhattan plots or even the presence or absence of certain peaks. In addition to confirming the relevance of *QTLs* detected in common using two reference genomes that represent the genetic structure of our panel, we identified at least three cases where our approach improves upon single-reference genome GWAS, and can thus be considered a promising alternative to pangenomic studies. **First**, we identified a unique case with *QTL* 7-1-LA1670 [**Fig. 5B]**: this *QTL* was detected only in the LA1670 reference genome because SNP calling failed to provide variants in the homologous region of LA1246. Interestingly, BLAST analyses revealed that the gene underlying *QTL 7- 1-LA1670* possesses multiple copies in both reference genomes, probably making this region very difficult to call. Second, in homologous regions, the statistical tests performed using the GWAS model (1) were affected by minor differences in both the kinship and population structure matrices. These variations led to changes in the p-values of markers within peaks, sometimes preventing them from reaching the significance thresholds **[Fig. 5A] [Table 3].** Third, structural variations in non-homologous regions between the two reference genomes influenced *QTL* detection in two distinct ways. In a first case, a large indel present in one reference genome made variant calling in the corresponding region of the other genome impossible (*QTL IG8-1-LA1670*). We hypothesize that the SNPs in this area could be interpreted as presence-absence variants. Alternatively, structural variations modified local LD between markers, such as with an inversion (*QTL IG3-1-LA1246*) or a small indel (*QTL IG2-1-LA1670*). Since *QTL* detection relied on the local score method, which depends on local LD, these structural variations had an impact on the results.

Using this method, we identified time- and temperature-dependent *QTLs* **[Table 3]**, with no common *QTLs* detected between the two temperature treatments, corroborating previous results observed in *A. thaliana* (22). The underlying QDR mechanisms induced under heat stress remain poorly understood. However, increasing evidence suggests that components of PTI and ETI are interconnected and characterized by different temporal induction dynamics (80). Studies on the *A. thaliana*/*P. syringae* pv. *tomato* DC3000 pathosystem has shown that a lower temperature favors ETI, while a higher temperature favors PTI (81). Moreover, the progression of the disease follows a temporal sequence in which the bacteria first invade the plant roots before colonizing the vascular system, with each stage involving different factors from both partners in the interaction (82). Our results support this observation, as a significant proportion of "strict candidate gene" orthologs were found to be expressed in the roots. This suggests that the defense mechanisms involved in our study may be dependent on time, temperature, and/or the specific plant organ. Given this complexity, we opted for a model using phenotypic ratings by Dai over a model that integrates temporal dynamics through measures, such as e.g. the ones used by Crispim and colleagues to decipher the genetic bases of beef cattle growth (83).

In the analyses of quantitative responses associated with bipartite plant-pathogen interactions, QDR genes identified were not only classical immunity genes: they have been described to be mostly associated with various pathways including detoxification, vesicle trafficking, epigenetic regulation, phosphorylation with kinase and associated receptors, phytohormone regulation and degradation of proteins and regulation of transcription (42,84). In our case, we decided to classify candidates by functional category. In addition, our analyses could allow, for the first time, definition of the nature of the genes underlying the *bwr-6* and *bwr-12 QTLs*, widely described over the last three decades as major *QTLs* conferring the QDR to *R. solanacearum* in tomato.

**The first functional category** corresponds to five *QTLs* for which the underlying homologs/orthologs, found either in the tomato ‘Heinz 1706’ cultivar or *A. thaliana*, have already been annotated as immunity genes in other pathosystems. Therefore, it suggests their involvement in broad-spectrum defense responses. Indeed, for *QTL 10-1-LA1246,* the orthologous underlying gene was found to have modulated expression in *A.thaliana* under attack by the cabbage leaf curl virus (CaLCuV) (85). According to TAIR, it is involved in « xenobiotic detoxification by transmembrane export across the plasma membrane». Micro- array data obtained from *A. thaliana* challenged by the *Pseudomonas syringae* pv *tomato* DC3000 strain led to the identification of the ortholog of *QTL 4-1-LA1246* (UDP- glucuronosyl/UDP-glucosyltransferase) (86). Interestingly the GDSL esterase/lipase (*QTL 9- 1-BOTH*) ortholog of *A.thaliana* is known to be down-regulated by abscisic acid treatment, leading to lower resistance towards the pathogen *Pseudomonas syringae* pv. *tomato* strain 1065 (87). *QTL 1-4-BOTH* corresponding to the *Solyc01g105270* ‘Heinz 1706’ cultivar ortholog coding for SlTOM1b (*S.lycopersicum* TOBAMOMIVUS MULTIPLICATION PROTEIN 1b), was annotated as an immunity gene involved in tobamovirus defense responses (88). The most significant SNP falls inside an exon of this gene. Translation of the CDS into protein reveals that one allele codes for a Methionine (M) whereas the other codes for an Isoleucine (I). Interestingly, M-I are non-synonymous substitutions that have already been shown to affect protein function (89, 90). Moreover, since some TOM (TOBAMOVIRUS MULTIPLICATION) proteins have transmembrane domains (91), different alleles of the *SlTOM1b* gene may play a role in protein stability inside the plasma membrane. Finally, the *Solyc12g008960* ortholog underlying *QTL IG12-1-BOTH* was identified in a transcriptome analysis of tomato leaves infected by thrips transmitting the tomato spotted wilt orthotospovirus (TSWV) (92).

**The second functional category** is constituted of calcium-related candidate genes underlying *QTLs 3-1-BOTH* and *IG12-1-BOTH*. Calcium is a universal second messenger contributing to either multiple pathogen or temperature-dependent signaling. However, information on calcium concentration variation during combined heat/pathogen stress is lacking (93). The *Solyc03g123790* gene which encodes a putative calcium/proton exchanger (*QTL*-3-1-BOTH) could participate in such variation of the Ca*^2+^* flux and is thus an interesting candidate gene. Variation of Ca*^2+^* concentrations are then relayed by calcium sensors, such as Calmodulin (CaM), Calmodulin-like (CML), Calmodulin-binding (CBP) proteins and Calcium-dependent protein kinases (CDPK) leading to the deployment of stimuli-specific responses (93). One of these sensors could be the CBP/IQM1 encoded by the *Solyc12g008960* gene (*QTL IG12-1-BOTH*). Many pieces of evidence make good arguments for the pertinence of the candidate: the first is statistical and has been previously discussed: this gene was among the most differentially expressed genes found in response to a 72-hour-post-thrips inoculation of TSWV (92). Based on several BLAST analyses using genes located within the *bwr-12 QTL* (94), the location of *QTL IG12-1-BOTH* was found to be approximately 600kbp upstream of the supposed location of *bwr-12* on our reference genomes. Another argument in favor of this discovery is that *bwr-12* is specific to phylotype I strains to which GMI1000 belongs. Furthermore, the *bwr-12 QTL* was found to be thermosensitive, with a reduction of the *bwr- 12* effect during heat stress (32). However, the temperature was not precisely regulated because experiments were carried out in greenhouses (temperature variations between 32 and 38°C) with a mean temperature of up to 36.5°C (32). Thus, the heat stress was higher than the one applied in our experiments. As for molecular arguments, IQM1 is known to bind to Catalase 2 (*At4g35090*) (95) in *A. thaliana,* and a similar mechanism could have been pinpointed in this study since two catalases were identified within *QTL 1-3-LA1246*. Despite the orthology between *Solyc01g100630* and *Solyc01g100640* in *A.thaliana* not having been clearly identified **[Table 3]**, Catalase 2 had the highest BLAST score.

**The third functional category** of candidate genes is constituted of receptor kinases. Indeed, to trigger ETI, plants have to recognize effectors that are molecules secreted by pathogens and they use many receptor-like kinases located on the plasma membrane. The recognition mechanism is based on the binding of effectors, which trigger their phosphorylation leading to a transduction signal in plant cells involving numerous other pathways forming ETI (42). Genes underlying *QTL 4-2-BOTH* and *QTL IG6-1-BOTH* correspond to receptor kinases. Using previously obtained fine mapping results (96); the position of *IG6-1-BOTH* was likely to be inside *QTL bwr-6*. Interestingly, *bwr-6* confers resistance to both phylotype I & II strains (32).

**The fourth class** of identified candidate genes is made up of genes belonging to gene families known to be involved in resistance against some pathogens. However, for this class, neither tomato nor *A.thaliana* orthologs have been described as taking part in responses to biotic stress. Sumo activating enzyme 1b Molybdenum cofactor biosynthesis (*QTL 3-4- LA1670*) is involved in the ubiquitination process (97), leading to protein degradation. The *QTL 1-1-LA1670* gene codes for a Subtilisin-like serine protease, a protein family that plays a role in pathogen recognition and immune priming (98). Finally, some glutathione S-transferases have been demonstrated to confer resistance to diseases, such as Fusarium head blight, using their detoxification ability (99). This could be also the case in our study for the gene underlying *QTL 9-2-BOTH*.

**The final class** contains all remaining genes. Sometimes no information about them is available or their link with immunity is not straightforward. However, previously cited reviews on QDR have shown that QDR involves various pathways, some of which have probably not yet been discovered (42, 84); this is all the more possible since each specific combination of stresses leads to a specific plant response (37).

Overall, by examining the natural diversity of responses to two combined stresses within a specific collection of wild tomato species accessions, our study confirms the significant impact of elevated temperatures on the defense mechanisms activated against *R. solanacearum*. This panel proves to be a valuable resource for identifying new thermostable resistance mechanisms. The development and characterization of a unique genomic resource, along with its use with two internal reference genomes in GWAS, have enabled the identification of several promising candidate genes, both common and specific to the two reference genomes. The next phase will involve functionally validating the genes involved in responses to combined stresses and evaluating their protective spectrum, as they could potentially represent a robust source of resistance based on their annotations.

## Data Availability Statement

The two reference genomes sequenced and annotated in this study have been deposited in the GenBank database under the accession numbers JASTWH000000000 (LA1246) and JASTWI000000000 (LA1670). The initial polymorphism matrices, derived from panel sequencing and SNP calling based on LA1246 and LA1670 reference genomes, are available *via* the following DOIs: https://dx.doi.org/10.25794/published_result/mk3v99ws (based on LA1246) and https://dx.doi.org/10.25794/published_result/bh8cvjct (based on LA1670). All phentotypic and genotipic as well as scripts developed and used in this study were archived in a publicly accessible platform at https://forgemia.inra.fr/adrien.belny1/gwas_belny_et_al_2025/.

## Author Contributions

R.B. conceptualized the project. R.B. and F.V. acquired funding for its achievement.

R.B., T.M.H supervised the project. G.B. and C.L. provide wild tomatoes accessions collection and enabled its sequencing and SNP calling. S.C. and J. G. analyzed sequence data, assembled, annotated and build a genome browser for the two reference genomes. H.D. and R.B. designed experiments. M.R. and H.D set-up and performed phenotyping experiments. H.D. analyzed the phenotypic traits. A.B., H.D. performed structure and LD analyses. A.B., H.D. created models, performed the GWA mapping and the genome-wide local score analysis. F.R., T.H.M., A.E., L.G. and R.B. verified experiments and analyses outputs. A.B, H.D. and R.B. wrote the initial draft of the manuscript. All authors contributed to critical discussions and the revisions.

## Acknowledgments

This project was funded in the framework of the INRAE / SYNGENTA BURNED/Rethink bilateral research project and subsequent addendum (N°15000441). We acknowledge SYNGENTA for funding the PhDs of H. Desaint and A. Belny and for technical support; Ludovic Legrand for his help with bioinformatic tools; and Caitlin Griffiths and Rebecca Stevens for reading and correcting the English usage.

## Supporting information captions

**S1**: Information on panel composition.

**S2**: Sequencing data for reference genomes and the mean coverage by SNPs for all accessions of the panel with raw data after SNP calling using the two reference genomes.

**S3**: Supplementary information on sequencing methodology and assembly.

**S4**: Part of variance explained by PCA axes for both reference genomes. **A:** LA1246. **B**: LA1670. A slight decrease is observed when the number of axes is above three.**S5**: Heritability (h^2^) calculated per Dai for experiments performed at 28°C and 32°C.

**S5:** Heritability (h^2^) calculated per Dai for experiments performed at 28°C and 32°C. Blue bars indicate heritability for the 28°C experiment and orange bars for the 32°C one.

**S6**: Average climate data recorded monthly between 1990 and 2019, relative to the GPS coordinates of LA1246 (**A**: Loja; Ecuador; 3,99S 79,36W; elevation 1259m, Köppen climate classification: Cfb) and LA1670 (**B**: Sama Grande; Tacna; Peru; 17,833S 70,51W; elevation 452m; Köppen climate classification: Bwh). The red line corresponds to the average temperature recorded while the average amount of precipitation is represented by blue histograms. [https://climatecharts.net/] [Zepner et al., 2021].

**S7:** Fixation index (F ST) within identified clusters and population structure of the panel analysed based on the LA1670 reference genome. **A**: F ST was computed using vcftools (using a window size of 10kb on the GWAS SNP matrix). Then the weighted means (by the number of markers within these windows) was computed between all combinations of the two clusters. **B**: Kinship matrix and representation of population structure: individuals were ordered following their clustering based on PCA axes to let blocks corresponding to structure appear. Results are consistent between the two figures: high kinship coefficients between individuals of two different clusters lead to small FST values, indicating clusters have similar genetic diversity (for example clusters 4-5).

**S8**: Evolution of linkage disequilibrium (LD; *r^2^*) along chromosomes for both reference genomes. **A**: LA1246; **B**: LA1670. X-axis: positions on chromosomes are indicated in base pairs. Order of magnitude of half-LD is a few kb. Except for chromosomes 5, 6 and 8.

**S9**: Half-decay in base pairs for all chromosomes for both reference genomes.

**S10:** LD (*r^2^*) between pairs of SNPs. For chromosome 1 (**A**-**B**) and 8 (**C**-**D**). Raw LD is represented on the left (**A**-**B**) and LD corrected with kinship on the right (**B**-**D**). Small blocks of SNPs in LD appear on chromosome 1. However, large blocks appear on chromosome 8, which is consistent with previous results on the LD half-decay.

**S11:** Manhattan plots, their related local score and QQ-plots. For each panel, data on the left are for the LA1246 reference genome while data on the right are for LA1670. Manhattan plots with local score (*ξ* = 3) are shown at the top of each panel. Red and blue lines represent the maximal and the minimal significance threshold computed per chromosome with autocorrelation between SNPs. In the middle of each panel are the Manhattan plots (with p- values; the dotted line represents the Bonferroni threshold) and the corresponding QQ plots (with p-values) are at the bottom of the figure. Each panel corresponds to a temperature condition and a Dai: **A**: 28°C-Dai7; **B**: 28°C-Dai10; **C**: 32°C-Dai4; **D**: 32°C-Dai5; **E**: 32°C- Dai6; **F**: 32°C-Dai7; **G**: 32°C-Dai10.

